# Novel regulation from novel interactions: Identification of an RNA sponge that controls the levels, processing and efficacy of the RoxS riboregulator of central metabolism in *Bacillus subtilis*

**DOI:** 10.1101/814905

**Authors:** Sylvain Durand, Adam Callan-Sidat, Josie McKeown, Stephen Li, Gergana Kostova, Juan R. Hernandez-Fernaud, Mohammad Tauqeer Alam, Andrew Millard, Chrystala Constantinidou, Ciarán Condon, Emma L. Denham

## Abstract

Small RNAs (sRNAs) are a taxonomically-restricted but transcriptomically-abundant class of post-transcriptional regulators. While potentially of importance, we know the function of few. This is in no small part because we lack global-scale methodology enabling target identification, this being especially acute in species without known RNA meeting point proteins (e.g. Hfq). We apply a combination of psoralen RNA cross-linking and Illumina-sequencing to identify RNA-RNA interacting pairs *in vivo* in *Bacillus subtilis*, resolving previously well-described interactants. Although sRNA-sRNA pairings are rare (compared with sRNA/mRNA), we identify a robust example involving the unusually conserved sRNA (RoxS/RsaE) and an unstudied sRNA that we term Regulator of small RNA A (RosA). This interaction is found in independent samples across multiple conditions. Given the possibility of a novel associated regulatory mechanism, and the rarity of well-characterised bacterial sRNA-sRNA interactions, we mechanistically dissect RosA and its interactants. RosA we show to be a sponge RNA, the first to be described in a Gram-positive bacterium. RosA interacts with at least two sRNAs, RoxS and FsrA. Unexpectedly, it acts differently on each. As expected of a sponge RNA, FsrA is sequestered by RosA. The RosA/RoxS interaction is more complex affecting not only the level of RoxS but also its processing and efficacy. Importantly, RosA provides the condition-dependent intermediary between CcpA, the key regulator of carbon metabolism, and RoxS. This not only provides evidence for a novel, and functionally important, regulatory mechanism, but in addition, provides the missing link between transcriptional and post-transcriptional regulation of central metabolism.

## INTRODUCTION

To adapt to changing environments and survive exposure to harsh conditions, organisms have evolved complicated metabolic and genetic regulatory networks to ensure that a homeostatic balance is maintained ^1, 2^. At the RNA synthesis level, gene expression can be modulated through combinations of transcription factors controlling genes required for growth and survival under specific conditions ^3–5^. At the post-transcriptional level, small regulatory RNAs (sRNAs) act to temper gene expression by short imperfect base pairing with their mRNA targets, altering the level of protein production by increasing or decreasing access to the ribosome-binding site, or by facilitating or blocking the access to the mRNA by ribonucleases (RNases) ^6, 7^. Most regulatory RNAs are independently expressed under the control of specific transcription factors. However, more recently, it has been shown that sRNAs can also be produced by processing RNAs that have other functions in the cell, such as tRNAs ^8^ and mRNAs ^9^.

Regulation by RNA is an important mechanism for fine-tuning gene expression in the Gram-positive model bacterium *Bacillus subtilis*, recently reviewed in ^10^. Over 150 potential sRNAs have been identified in *B. subtilis* and shown to be expressed in a condition-dependent fashion ^11–13^. To date the roles of very few of these putative sRNAs have been determined. However, where targets have been identified, they have been shown to play key roles in stress adaptation. *B. subtilis* notably expresses three sRNAs with C-rich regions (CRRs) with similar predicted secondary structure; RoxS/S415 (Related to oxidative stress) ^14^, FsrA/S512 (Fur regulated small RNA) ^15^ and CsfG/S547 (Controlled by sigma-F and sigma-G) ^16^, (S numbers relate to transcriptionally active segments identified by Nicolas *et al.* ^12^). The RoxS sRNA is one of the best characterised sRNAs in Gram-positive bacteria ^14, 17, 18^ and is conserved among Bacilli and Staphylococci, where it is named RsaE ^19, 20^. RoxS has been shown to be upregulated in response to nitric oxide (NO) in *B. subtilis* and *S. aureus*, by the two component system ResDE, and its homolog SsrAB, respectively ^14^. RoxS expression is also activated when malate is supplied as a carbon source. This control is mediated by the transcription factor Rex, that is known to sense the NAD/NADH ratio of the cell. Indeed, this ratio is perturbed by the conversion of malate to pyruvate by the three malate dehydrogenases of *B. subtilis* that reduce NAD^+^ to NADH, and by its cycling through the TCA pathway. It has been proposed that one role of RoxS is to re-equilibrate the NAD/NADH ratio of the cell by inhibiting the expression of enzymes leading to the production of NADH. FsrA is regulated by the transcription factor Fur and acts as part of the iron-sparing response ^15^. Fur down-regulates mRNAs whose protein products contain iron as part of their structures, but are not essential for growth, therefore ensuring iron availability for essential iron-containing proteins ^15, 21^. Interestingly, both RoxS and FsrA down-regulate several genes encoding enzymes of the TCA cycle that produce NADH. CsfG is highly expressed during sporulation, anaerobic growth and after glucose exhaustion ^12^. During sporulation, expression of this sRNA is controlled by the sigma factors F and G which are restricted to the forespore ^16^. However, to date no mRNA target or physiological role for CsfG has been identified. Durand *et al.* have hypothesised that its similar sequence motifs and structure to RoxS and FsrA suggests these three sRNAs may have overlapping targets and play similar roles under different growth conditions ^14^.

The lack of well resolved pathways through which sRNAs act in no small parts reflects the difficulty of global scale target identification, this being more acute in some bacteria than others. In many enterobacteria, such as *Escherichia coli* and *Salmonella typhimurium*, the RNA chaperone Hfq plays a key role as a mediator of sRNA-mRNA interactions and has greatly enabled the identification of mRNA targets through pull-down studies ^22, 23^. Although Hfq is conserved in Gram-positive bacteria, it does not appear to play a global role in RNA-mediated regulation of gene expression ^24, 25^. Hfq-dependent regulation by only one sRNA in *Listeria* and a handful in *Clostridium* are the only known exceptions. It is therefore generally accepted that sRNA regulation in the Firmicutes either depends on different RNA chaperones or can occur in the absence of any protein factors. A number of groups have used *in vivo* RNA cross-linking with the psoralen AMT, followed by ligation to form chimeras and RNAseq to identify RNA-RNA interactions in eukaryotic cells ^26–28^. Here then we employed LIGR-seq ^26^ to identify sRNA targets in *B. subtilis*. In addition to identifying many known members of the FsrA and RoxS regulons and several new targets, we also identified a new regulatory RNA, S345, that interacts with both FsrA and RoxS. These interactions are found in independent samples and across multiple conditions. Given the possibility of a novel associated regulatory mechanism, and the rarity of well-characterised bacterial sRNA-sRNA interactions, we mechanistically dissect S345 and its interactants. We show that S345 not only functions as an RNA sponge for RoxS, but also affects its processing and degradation. We rename this sRNA RosA (for <u>R</u>egulator <u>o</u>f <u>s</u>RNA <u>A</u>). We show that the transcription of RosA is under the control of the carbon catabolite control protein A (CcpA), linking the action of RoxS to the carbon source availability in *B. subtilis*.

## MATERIALS AND METHODS

### Media and growth conditions

Selection for transformations was performed on Lysogeny Broth (LB) at 37°C supplemented with required antibiotics. For *E. coli* these were ampicillin (100 μg ml^−1^) or chloramphenicol (10 μg ml^−1^) and for *B. subtilis* either phleomycin (4 μg ml^−1^), kanamycin (20 μg ml^−1^), tetracycline (5 μg ml^−1^), chloramphenicol (5 μg ml^−1^), erythromycin (2 μg ml^−1^), spectinomycin (100 μg ml^−1^) or combinations of the above. Growth experiments were performed in LB, M9 medium supplemented with glucose at a final concentration of 0.3% ^12^ or MD medium ^29^ supplemented with arabinose or malate at a final concentration of 1%.

### Bacterial strain construction

All *E. coli* and *B. subtilis* strains and plasmids used in this study are listed in Supplementary Table I. Primer sequences can be found in Supplementary Table II. *E. coli* DH5α and TG1 were used for all cloning procedures. *B. subtilis* strains were derived from the *B. subtilis* 168 trp^+^. The isogenic deletion mutants were constructed according to the method described by Tanaka et al ^30^ without pop-out of the deletion cassette. Transfer of genetic mutations between strains was achieved by transformation of genomic DNA extracted from the relevant strain. Reintroduction of sRNAs under the control of their native promoters was achieved by Gibson Assembly into pRMC that integrates into the *amyE* locus ^31^ of a PCR amplicon. Primer annealing sites were chosen to include the native promoter mapped in Nicolas et al ^12^. The sequence of cloned sRNAs was subsequently confirmed by sequencing and transformed into *B. subtilis* (plasmid pRMC+P_native_-sRNA). Integration into the *amyE* locus was confirmed by an iodine halo assay by replica plating transformation plates onto starch plates. The RosA promoter fusion was constructed at the native genomic locus by integration of the pBSBII plasmid ^32^. Combinatorial strains were constructed in the genetic background of the same promoter fusion strain by transformation of genomic DNA of the respective strain and selection on the appropriate antibiotics.

### *In vivo* RNA interactome

#### AMT *in vivo* cross linking

Bacteria were grown to the required O.D before 10 O.D _600 nm_ units were harvested by centrifugation (4000 g, 5 minutes, 4°C). Bacteria were resuspended in 2 ml PBS either containing no AMT (to identify background and levels of spurious interactions) or 0.7 mM AMT. Bacteria were incubated for 10 minutes at 37°C for 10 minutes before being transferred to a 6 well plate. The bacteria were exposed to UV 365 nm at 0.120 Jcm^−2^ for 10 minutes before being added to 1 ml of ice cold killing buffer (20 mM Tris-HCl [pH 7.5], 5 mM MgCl_2_, 20 mM Na-azide). The bacteria were harvested by centrifugation at (4000 g, 5 minutes, 4°C). the supernatant was discarded and the pellet flash frozen in liquid nitrogen. We determined the *in vivo* RNA interactome of *B. subtilis* grown in M9 minimal media supplemented with 0.3% glucose at three points in the growth curve (exponential phase O.D._600nm_ 0.5, stationary phase O.D._600nm_ 1.4 and just after lysis had started to occur, and in LB at mid-exponential phase (O.D._600nm_ of 1.0). A *Δfur* mutant ^33^ was prepared in LB at mid-exponential phase to increase expression of the Fur regulated sRNA FsrA. Samples were prepared in duplicate.

#### RNA extraction and formation of chimeras between interacting RNAs

The RNA was extracted by resuspending the cell pellet in 800 µl LETS buffer (10 mM Tris-HCl [pH 8.0], 50 mM LiCl, 10 mM EDTA, 1% sodium dodecyl sulfate [SDS]) and bead beating in a FastPrep using 0.1 µm glass beads for three rounds of 40 seconds. The tubes were transferred to ice in between cycles. The tubes were briefly spun to remove the bubbles created during bead beating. Two rounds of phenol chloroform isoamyl alcohol extraction and one round of choloroform isoamyl alcohol extraction were carried out. Before the addition of 10 % v/v NaAcetate and 1 ml isopropanyl and precipitation of RNA overnight at −20°C. The RNA was pelleted by centrifugation at maximum speed at 4°C and the pellet was washed with 70% Ethanol before being air dried and resuspended in water. The RNA was quantified using the Qubit kit (Fisher Life Science). 10 µg of RNA was treated with Turbo DNase (Fisher Scientific) to remove contaminating DNA. Ribosomal RNA was removed using Ribozero (Illumina) according to the manufacturer’s instructions. To form the chimeric RNAs between RNAs crosslinked with AMT the protocol described by Sharma *et al.* was followed as described in the supplementary data ^26^. The only modification was the use of CircDNAligase (Epicentre) instead of CircRNAligase as this has been discontinued.

#### RNAseq

Following uncrosslinking at UV 254 nm, RNA was purified and resuspended in 10 µl H_2_O and processed through the TruSeq stranded total RNA library kit (150 bp) (Illumina) according to the manufacturer’s instructions. The resulting libraries were sequenced on the MiSeq (Illumina).

#### Analysis

STAR aligner was used to map reads (Version STAR_2.6.0c_08-11) (34). This mapping tool is designed to analyse splicing of introns and exons, which is similar to what is created through the formation of chimeric reads where two different RNA fragments have been joined together. By identification of reads that map to different features (protein coding sequences, sRNAs, UTRs, transcripts for ncRNAs such as rRNA and tRNA, or transcribed intergenic regions) it is possible to identify RNA interactions. STAR aligner was set to single end read mode to map read 1 and read 2 separately, the chimeric detection mode activated, as this has been reported to be more sensitive to chimeric junctions. The allowed mismatches in mapping was set to default for STARaligner. The output from STAR was merged in to one Sam file, before being annotated using featureCounts within the package subread-1.6.3 in R, with all further statistical analysis also carried out in R (35). In our initial analysis we found many reads mapped to the genome, but to unannotated features. To overcome this problem, we created new features for the unannotated regions of the genome and these are termed UA-start – stop in the data files.

Chimeric reads will map to two different genomic features, whereas non-chimeric reads should only map to one feature. The exception is of those reads with repetitive mapping or those that map to two neighbouring features such as genes in an operon. An interaction count table was generated of reads that mapped to more than one feature and thus are considered as interacting pairs. The interaction count matrix then allowed statistical analysis of each interacting pair using a hypergeometric test. This compares the number of interaction read counts for each specific interaction with the number of other interaction read counts formed by each member of that pair, but with other RNAs. All interaction pairs with a P-value below 0.05 were extracted as significant interactions. To increase confidence in the identified interactions a second analysis was carried out where the pairs of sequenced samples were analysed together and untranslated regions were combined with coding sequences. The hypergeometric test was repeated and a P-adjusted value was calculated using Benjamini and Hochberg to control for the false discovery rate which was set at 0.05 ^34^.

To add further confidence to which interacting pairs form the most likely interactions, the interacting pairs were further assessed by *in silico* prediction with IntaRNA2.0, which predicts the stability and binding position between two interacting RNA pairs (36,37). The gene and any untranslated region that has been identified associated with the gene of interest (12) were included in the prediction to take into account the transcriptional start and stop sites. If no UTR had been identified for an mRNA the 50 bp up and downstream of the start and stop site were employed.

#### Proteomics analysis

Strains were grown to O.D._600 nm_ 1.0 in LB. 20 O.D. units were harvested and washed 3 X with PBS to remove media components. Cells were resuspended in 200 µl urea buffer (8 M Urea, 50 mM Tris and 75 mM NaCl). 200 µl of urea buffer washed 0.1 µM beads were added to the cells before being disrupted using three rounds of bead beating for 40 seconds using a FastPrep. Cells were placed on ice between the three rounds of bead beating. The disrupted cells were then sonicated in a water bath for 15 minutes. Cell extracts were centrifuged at 15,000 × g, 5 min and supernatants used for protein quantification (Qubit protein assay kit). Protein reduction and alkylation was conducted by mixing 150 µg of total protein with 10 mM TCEP and 40 mM CAA, at 600 rpm, for 20 min at room temperature. After, proteins were predigested with 1.5 µg of rLysC (Promega) for 3 h at room temperature and samples diluted with 50 mM ammonium bicarbonate, 2 M urea final concentration. Protein digestion was performed with 1.5 µg of Trypsin (Promega) overnight at room temperature. The reaction was stopped by adding 1% TFA and 10 µg of peptides were desalted using StageTip ^35^.

Reversed phase chromatography was used to separate 1 µg of tryptic peptides prior to mass spectrometric analysis. The cell proteomes were analysed with two columns, an Acclaim PepMap µ-precolumn cartridge 300 µm i.d. × 5 mm, 5 μm, 100 Å and an Acclaim PepMap RSLC 75 µm i.d. × 50 cm, 2 µm, 100 Å (Thermo Scientific). The columns were installed on an Ultimate 3000 RSLCnano system (Dionex) at 40°C. Mobile phase buffer A was composed of 0.1% formic acid and mobile phase B was composed of acetonitrile containing 0.1% formic acid. Samples were loaded onto the µ-precolumn equilibrated in 2% aqueous acetonitrile containing 0.1% trifluoroacetic acid for 8 min at 10 µL min-1 after which peptides were eluted onto the analytical column at 250 nL min-1 by increasing the mobile phase B concentration from 8% B to 25% over 90 min, then to 35% B over 12 min, followed by a 3 min wash at 90% B and a 15 min re-equilibration at 4% B.

Eluting peptides were converted to gas-phase ions by means of electrospray ionization and analysed on a Thermo Orbitrap Fusion (Thermo Scientific). Survey scans of peptide precursors from 375 to 1500 m/z were performed at 120K resolution (at 200 m/z) with a 2×105 ion count target. The maximum injection time was set to 150 ms. Tandem MS was performed by isolation at 1.2 Th using the quadrupole, HCD fragmentation with normalized collision energy of 33, and rapid scan MS analysis in the ion trap. The MS2 ion count target was set to 3×103 and maximum injection time was 200 ms. Precursors with charge state 2–6 were selected and sampled for MS2. The dynamic exclusion duration was set to 60 s with a 10 ppm tolerance around the selected precursor and its isotopes. Monoisotopic precursor selection was turned on and instrument was run in top speed mode.

Thermo-Scientific raw files were analysed using MaxQuant software v1.6.0.16 ^35^ against the UniProtKB *B. subtilis* database (UP000001570, 4,260 entries). Peptide sequences were assigned to MS/MS spectra using the following parameters: cysteine carbamidomethylation as a fixed modification and protein N-terminal acetylation and methionine oxidations as variable modifications. The FDR was set to 0.01 for both proteins and peptides with a minimum length of 7 amino acids and was determined by searching a reversed database. Enzyme specificity was trypsin with a maximum of two missed cleavages. Peptide identification was performed with an initial precursor mass deviation of 7 ppm and a fragment mass deviation of 20 ppm. The MaxQuant feature ‘match between runs’ was enabled. Label-free protein quantification (LFQ) was done with a minimum ratio count of 2. Data processing was performed using the Perseus module of MaxQuant v1.6.0.16 ^36^. Proteins identified by the reverse, contaminant and only by site hits were discarded. Only protein groups identified with at least two assigned peptides were accepted and LFQ intensities were log2 transformed. Significantly regulated proteins were identified in two rounds of analysis. First, a Student’s T-test (FDR 0.05) and a minimum difference of S0=0.1 was applied on all biological replicates. Second, a finest statistical analysis was applied using the same parameters as before but removing the outliers identified by principal component analysis and Pearson correlation test. The significantly regulated proteins were selected from both analyses.

#### Plate reader experiments

Experiments to monitor promoter activity were carried out in a 96-well format in a BioTek Synergy Plate reader and analysed as described previously ^31^.

#### RNA isolation and Northern Blotting

RNA was isolated from mid-log phase *B. subtilis* cells growing in the indicated medium by the RNAsnap method described in Stead *et al.*, 2012. Northern blots were performed as described previously (Durand *et al.*, 2012). The S345/RosA riboprobe was transcribed *in vitro* using T7 RNA polymerase (Promega) and labelled with [a-^32^P]-UTP using a PCR fragment amplified with oligo pair CC2440/CC2441 as template. The oligos CC089, CC964 and CC875 were 5’ end-labelled with T4 polynucleotide kinase (PNK) and [g-^32^P]-ATP and used to probe *sucC, ppnkB* and RoxS RNA respectively.

#### Quantitation of sRNAs

S345/RosA and RoxS RNAs where transcribed *in vitro* from PCR fragments amplified with the oligo pairs CC2406/CC2407 and CC1832/CC1833 respectively. Known quantities (in fmol) of *in vitro* transcribed S345/RosA and RoxS RNAs, and 5 *μ*g total RNA isolated from wild-type cells were loaded on a denaturing 6% acrylamide gel. The oligos CC2347 and CC875 were 5’ end-labelled with T4 polynucleotide kinase (PNK) and [g-^32^P]-ATP and used to probe on Northern blot S345/RosA and RoxS respectively.

#### Electrophoretic mobility shift assays (EMSA)

For EMSA assays, S345/RosA, RoxS and FsrA sRNAs where transcribed with T7 RNA polymerase *in vitro* from PCR fragments amplified with the oligo pairs CC2406/CC2407, CC1832/CC1833 and CC2492/CC2493 respectively. A 15 *μ*l reaction was prepared by mixing 2 pmol of S345/RosA RNA with an increased concentration of RoxS or FsrA RNA (1, 2, 3 and 4 pmol) in 1X the RNA binding Buffer (10 mM tris pH8; 50 mM NaCl; 50 mM KCl, 10 mM MgCl_2_). The Mix was heated for 3 min and cool down at room temperature for 10 min. After cooling, 10 *μ*l of glycerol (Stock solution 80%) was added and the RNA were loaded on a 6% non-denaturing polyacrylamide gel (Acry:bisacry – 37.5:1). RNA was transferred on to a Hybond N+ membrane and hybridized with the S345/RosA radiolabelled probe (CC2347).

#### Strain Competition experiment

Strains marked with appropriate antibiotics were combined at a 1:1 ratio, inoculated at a starting O.D._600_ nm and grown for 24 hours in LB. To confirm starting ratios at a 1:1 ratio colony counts were performed on the initial inoculum. At 24 hours cultures were serially diluted and plated on LB plates containing the relevant antibiotics to enable counting of each strain. Ratios of strains were calculated and Welch’s T test was used to determine significance. An average of three technical replicates each containing three biological replicates was carried for each combination of strains.

## RESULTS

### *in vivo* RNA crosslinking identifies known and unknown sRNA-RNA interactions

To identify new sRNA-mRNA interactions in *B. subtilis* we applied the LIGR-seq protocol ^26^ to *B. subtilis* cells growing in M9 minimal media supplemented with 0.3 % glucose (exponential and transition phase) or in LB (WT and Δ*fur* mutant at exponential phase). The Δfur mutant was included to increase the expression levels of the sRNA FsrA, the transcription of which is repressed by Fur. Cells were irradiated at 365 nm with the chemical crosslinker AMT (4’-aminomethyltrioxsalen). Biological replicates of each sample were prepared. RNAs were extracted, ligated, and non-crosslinked RNA was digested with RNase R. Crosslinks were reversed with 254 nm irradiation and RNA samples were subjected to high-throughput sequencing to detect chimeras formed by ligation.

We designed and analysed the resulting RNA-seq data for chimeras using a customized pipeline. This included using STAR aligner which is designed for mapping RNAseq data containing splicing of introns and exons in data sets produced from eukaryotes ^37^. We discovered that carrying out the alignment using single-end read mode and activating the chimeric detection increased the sensitivity of the chimeric read detection. This also enabled us to map chimeric reads where the ligation of the two fragments occurred close to the read ends.

In each of the eight individual samples analysed, many potential RNA-RNA interactions were identified through using the customized pipeline (see Methods). However, to validate the data, we focused on chimeras identified for the well-characterized sRNAs of *B. subtilis*, FsrA and RoxS (Supplementary Table 3 A (FsrA) and B (RoxS)). Many known interactions of FsrA such as *citB, gltAB, lutA* and *leuC* ^15, 21^ and for RoxS, *citZ* ^14^ were identified in our screen. However, many other interactions were also present in the data set. To improve our confidence in identifying new targets of FsrA and RoxS we combined the data from the sample pairs and reanalysed the data. Statistically significant interactions and the P-adjusted value for each sample pair are shown in Supplementary Table S4 A (FsrA) and B (RoxS). The IntaRNA prediction of each interaction is also shown. Several potential new targets that have a possible link with iron metabolism were identified for FsrA. For example, we identified many chimeras between FsrA and the *yydF* mRNA, encoding a secreted peptide that controls LiaRS activity ^38^. The gene downstream of *yydF* in this operon, *yydG*, encodes a protein that contains an Fe-S cluster and is part of a protein complex required to process YydF into a functional peptide. RoxS regulates the expression of many RNAs encoding proteins involved in central metabolism, such as citrate synthase (CitZ) ^14^. Our data showed a statistically significant interaction between RoxS and the *citZ* mRNA and also for *odhA* which encodes 2-oxoglutarate dehydrogenase (E1 subunit).

The above data confirm the validity of the LIGR-seq technique to identify new potential sRNA-mRNA interactions in bacteria. We suggest therefore that as a method AMT crosslinking may be considered as being complementary to studies focusing on individual RNAs such as MAPS ^39^ and those focusing on finding RNA interactions that occur on proteins such as RILseq ^23^ or CLASH ^40^, where different interacting RNAs have been found depending on the technique used.

### Identification of a novel robust sRNA-sRNA interaction

Analysis of the RNA interactome also allowed us to map sRNA-sRNA interactions. Indeed, the most statistically significant interaction for both FsrA and RoxS was with the predicted sRNA S345 and S346 (annotated as 3’ UTR of S345) (Figure 1). The interaction between S345 and both RoxS and FsrA was the most represented chimera pair in the interactions that we detected between RoxS and FsrA. The interaction is not only of strong statistical significance but was found in multiple growth conditions (Supplementary tables 3,4). As robustly described sRNA-sRNA interactions are unusual (for other examples see ^41, 42^) we sought to characterize this further.

**Figure 1.**
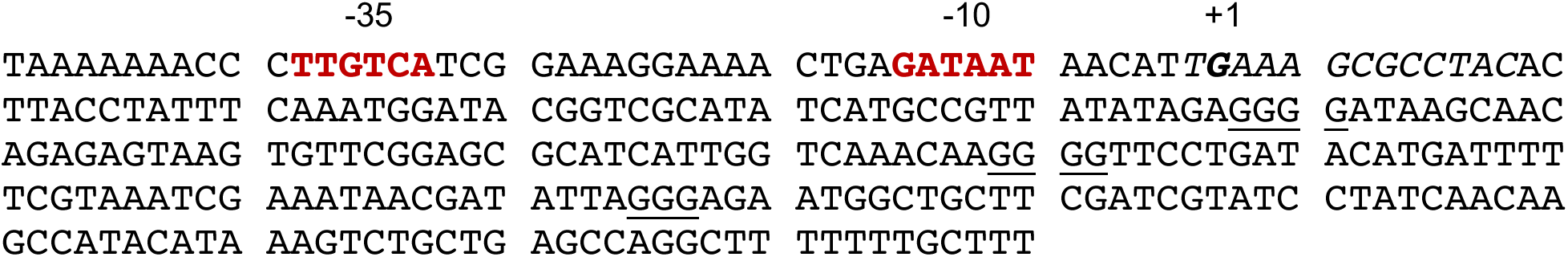
Sequence of S345 and its promoter region. The −10 and −35 sequence of the putative σA promoter are indicated in red. The Cre site for CcpA binding starting at −1 is italicized. The 3 G-rich regions (GRR) of S345 are underlined.

The sequence of S345 has three G rich regions (GRRs) (Figure 1 and 2A) with potential complementarity to the C-rich regions (CRRs) of FsrA and RoxS that have been shown to be involved in the interactions with their mRNA targets. We used RNAfold to predict how FsrA and RoxS might interact with S345 (Figure 2B and supplementary Figure 1) ^43^. The interaction with FsrA is predicted to incorporate GRR2 of S345 and CRR2 of FsrA (Supplementary Figure 1). Intriguingly, the interaction with RoxS is predicted to incorporate both GRR1 and GRR2 of S345, and CRR1, CRR2 and CRR3 of RoxS (Figure 2B). Both predictions include two long stretches of interacting nucleotides, suggesting these two RNA pairs can form stable duplexes.

**Figure 2.**
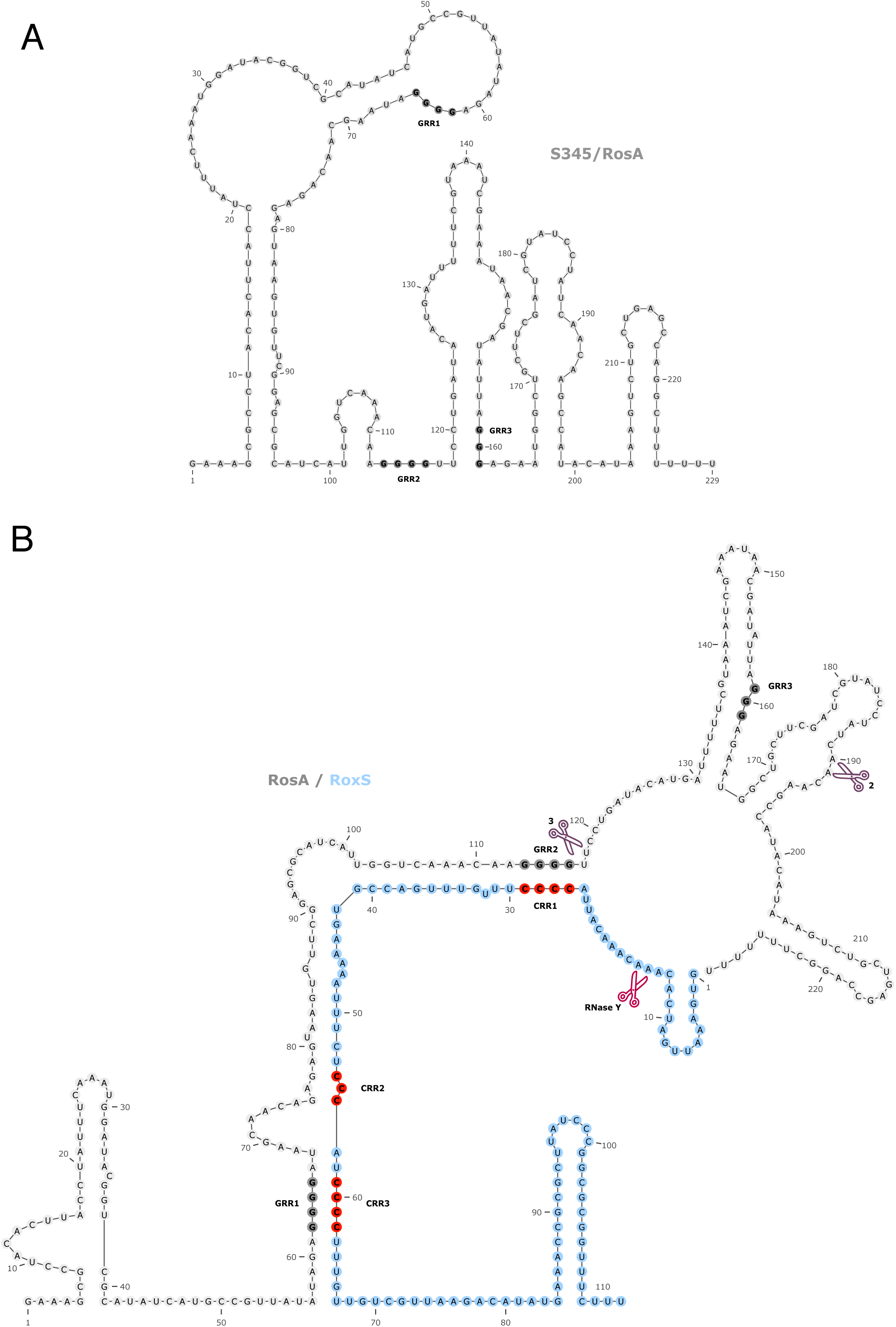
Prediction of the interaction between S345 and RoxS. **A.** Prediction of the secondary structure of S345 with RNAfold web server (http://rna.tbi.univie.ac.at/cgi-bin/RNAWebSuite/RNAfold.cgi) **B.** The interaction between RoxS and S345 sRNAs was predicted with IntaRNA web server (http://rna.informatik.uni-freiburg.de/IntaRNA/Input.jsp). The RoxS sRNA is coloured in blue. The C-rich region of RoxS and the G-rich region of S345 are coloured in red and grey respectively. The RNA processing site of RoxS and the two main processing site of S345 are indicated with red and purple pairs of scissors respectively (see also Fig. 5).

### RoxS interacts directly with S345 *in vitro*

To confirm the potential interaction between S345 and RoxS or FsrA, we performed an Electrophoretic Mobility Shift Assay (EMSA; Figure 3). S345 was mixed with increasing concentrations of RoxS or FsrA and loaded on a non-denaturing acrylamide gel. The results show that RoxS can bind very efficiently to S345, producing a sharp band of higher molecular weight and a full-shift of S345 even at the lowest molar ratio of RoxS to S345 tested (0.5). Complex formation between FsrA and S345 was less efficient and the complex was less well defined, but nonetheless visible. A full shift of S345 was not apparent even at a 2-fold excess of FsrA (Figure 3). When both RoxS and FsrA were incubated together with S345, the interaction was clearly in favour of RoxS, with only trace quantities of the FsrA-S345 complex visible. These results suggest that RoxS has a higher affinity for S345 than FsrA and are in agreement with the longer predicted duplex between these two sRNAs.

**Figure 3.**
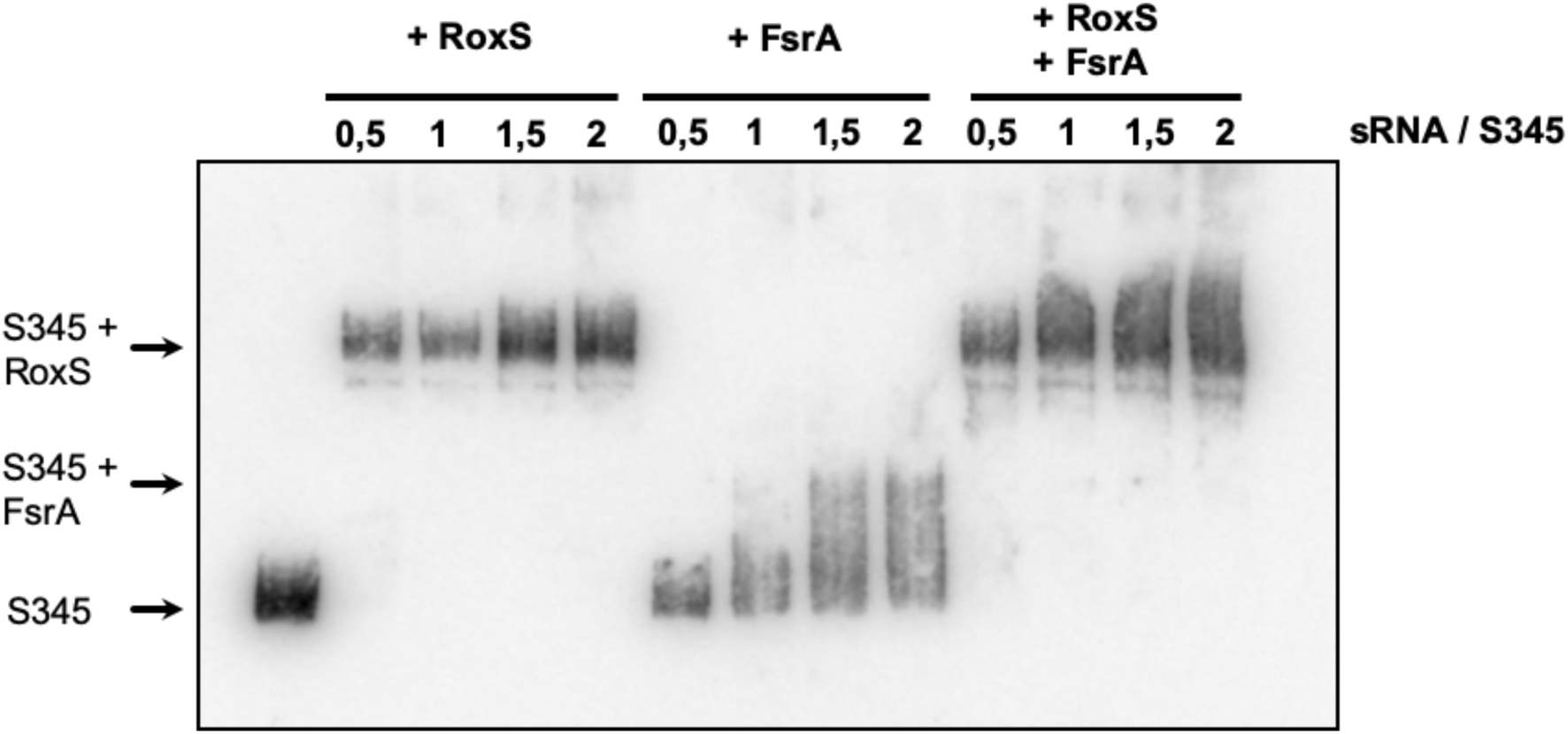
S345 interacts with RoxS and FsrA sRNAs. Electrophoretic mobility shift assays (EMSA) of S345 with RoxS and FsrA. 1 pmol of S345 was incubated with an increasing concentration of RoxS and/or FsrA. The ratio of sRNA/S345 is indicated.

### S345 is a highly processed sRNA

To begin to characterize S345, we first assayed its expression pattern and stability in the same conditions as those used in the crosslinking experiment (LB and in M9 minimal medium + glucose). Northern blot analysis of total RNA isolated at different times after the addition of rifampicin to block new transcription showed that the level of the S345 RNA is higher in LB than in M9 at mid-exponential phase (Figure 4). Moreover, three major forms of S345 were detected. The approximate sizes for species 1, 2 and 3 are 230 nts, 185 nts and 120 nts, respectively (Supplementary Figure 2A). The half-life of the largest species (1) was less than 1 minute, while the dominant species (2) had a slightly greater stability, with a half-life of 1.9 minutes in LB. The shortest species (3) had the longest half-life: 7.1 minutes in LB (Figure 4). Since the half-lives of the three forms of S345 are similar in M9 + glucose, the lower levels of S345 in this medium are most likely due to transcriptional regulation (see below).

**Figure 4.**
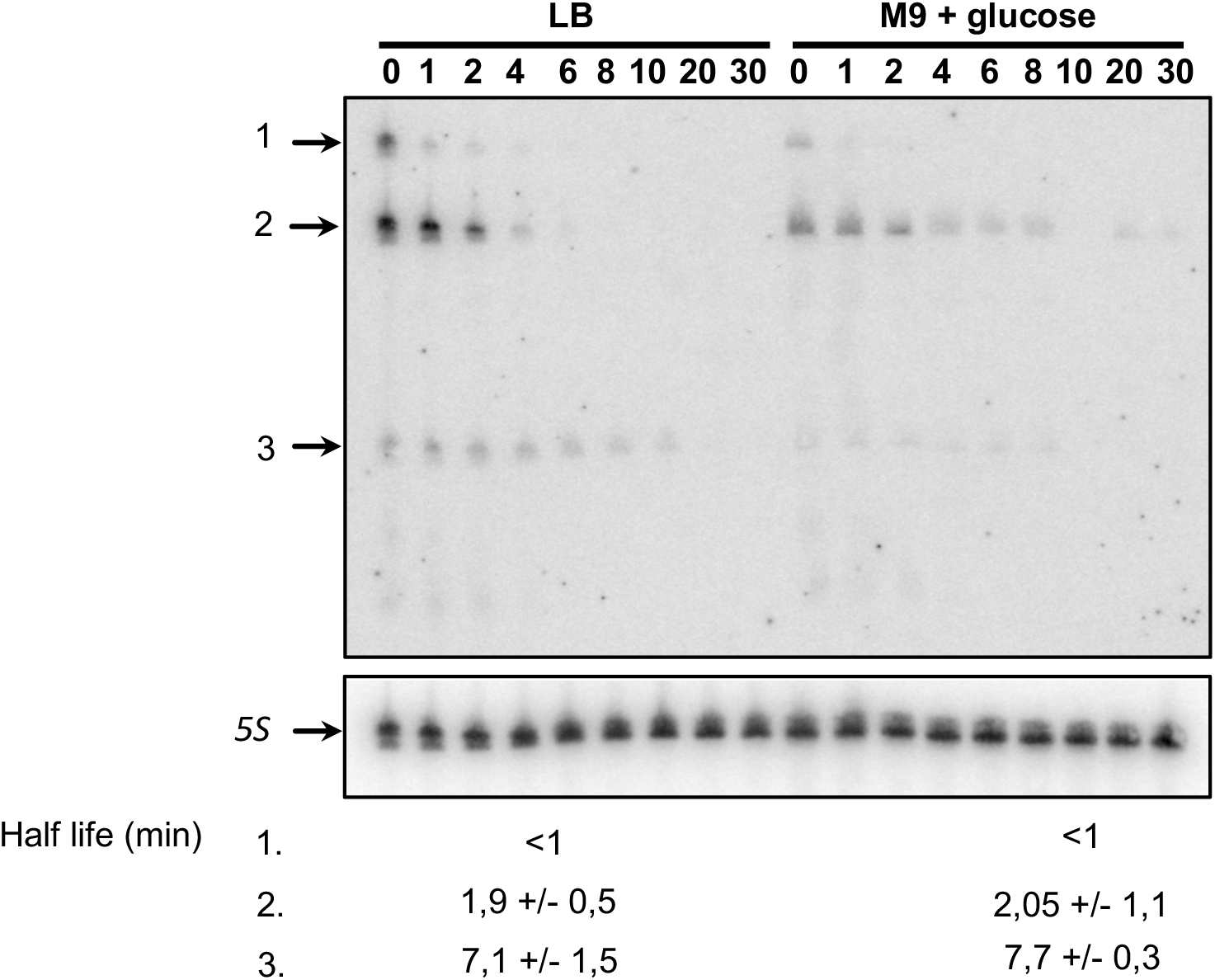
S345 is a differentially expressed, highly processed sRNA. A representative Northern blot showing total RNA isolated at times after addition of rifampicin (Rif) to WT strain grew in LB or M9 with glucose (0,3%). The three different forms of S345 are indicated by arrows. For each form of S345, the half-life with their standard deviation calculated from two independent experiments (biological replicates) are given under each autoradiogram.

The 5’-end of S345 was suggested from the sequencing product of the LIGR-seq data and was confirmed by primer extension using an oligo close to the putative S345 transcriptional terminator (supplementary Figure 2B). We were able to predict a putative sigma-A promoter that fits perfectly with this mapped 5’ end (Figure 1). Moreover, the distance between the mapped 5’ end and the putative transcriptional terminator is 229 nts, which corresponds well with the size of the largest band detected by Northern blot and suggests that species 1 corresponds to the primary S345 transcript.

The Northern blot in Figure 4 was performed with an oligonucleotide probe starting 30 nts from the 5’ end of S345. A second probe starting only 10 nts from the 5’ end gave a similar pattern (data not shown). We thus deduced that the three major forms of S345 have the same 5’ end and that species 2 and 3 are processed from the primary transcript at 3’ proximal sites. In agreement with this hypothesis, when S345 was first identified by tiling array, an extended 3’ region was identified that was annotated as S346 ^12^. The size of our proposed primary transcript corresponds to the sum of the annotated segments S345 + S346. Our LIGR-seq data showed numerous truncations of S345 at its 3’ end and allowed us to determine an approximate position for the cleavage site generating species 2 (Figure 2B). The processing of the 3’ end of S345 was further confirmed as we were also able to map the 5’ end of a 3’ degradation product (*) stabilized in a *ΔrnjA* mutant strain by primer extension. This corresponds to an endonucleolytic cleavage event occurring at the end of the duplex between RoxS and S345 (Supplementary Figure 2 and Figure 2B). The upstream cleavage product, protected from degradation due to its hybridisation with RoxS corresponds to the smallest (120 nts) S345 species (species 3). These observations suggest that S345 is quickly processed near its 3’ end to form species 2 and 3, in agreement with the shorter half-life of the full length S345 RNA compared to its two derivatives (Figure 4).

### S345 destabilises RoxS

To determine whether S345 had an effect on RoxS levels or stability *in vivo*, we measured the rate of RoxS RNA degradation before and after the addition of rifampicin to WT and ΔS345 mutant strains. The experiment was done in LB, since S345 is expressed at higher levels in this medium. Samples were taken over a time course of 0 to 60 minutes and the RNA analysed by Northern blot. RoxS expression was significantly higher in the absence of S345 (Figure 5A). The half-life of RoxS in the presence of S345 was 13.2 minutes, whereas in the absence of S345 the half-life increased to 46.3 minutes. This result shows that expression of S345 leads to destabilization of the RoxS sRNA. In contrast, deletion of S345 has no impact on the stability of FsrA in LB media (Supplementary Figure 3).

**Figure 5.**
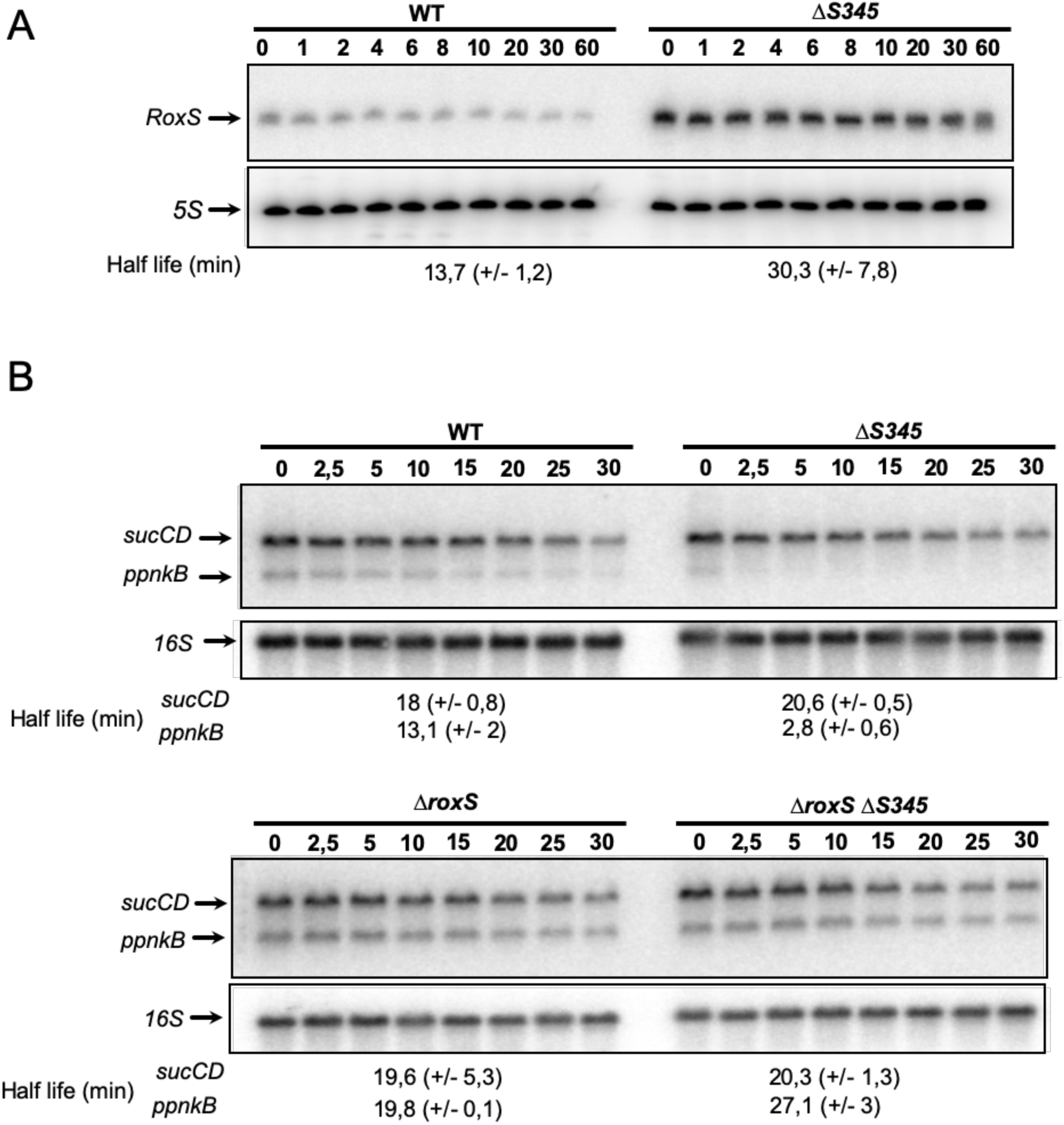
Deletion of S345 alters the turnover rate of RoxS and RoxS targets. **(A)** A representative Northern blot of total RNA isolated from wild-type (WT) and *ΔS345* strains probed for the RoxS sRNA at times after addition of rifampicin (Rif) at 150 μg/mL. The blot was re-probed for the 5S rRNA as a loading control **(B)** Northern blot of total RNA isolated from wild-type WT, *ΔS345*, *ΔroxS* and *ΔS345 ΔroxS* strains probed for the *sucCD* or *ppnKB* mRNA. The blot was re-probed for 16S rRNA as a loading control. Calculated half-lives are shown beneath the autoradiographs and are the average of 2 experiments, with standard errors as shown.

We also calculated the relative amount of S345 and RoxS present in the cells grown in LB. In 5 µg of total RNA, S345 and RoxS were present at approximately equimolar amounts (10 fmol each; supplementary Figure 4). This result shows that there is sufficient S345 in the cell to completely titrate all RoxS present in the cell under equilibrium conditions and suggests that it could act as an RNA sponge to counteract RoxS activity by titrating it away from its targets.

### Deletion of S345 leads to destabilisation of FsrA and RoxS targets

RoxS has been previously shown to negatively impact the stability of the *ppnKB* and *sucCD* mRNAs encoding an NAD(H) kinase and succinate dehydrogenase, respectively ^14^. If S345 indeed modulates the availability of RoxS to interact with its targets, we would predict that the half-life of these transcripts would decrease in the *ΔS345* strain due to the additional free RoxS in the cell (Figure 5A). In Northern blot experiments performed on cells growing in LB medium, the half-life of the *ppnKB* mRNA was indeed decreased 4.7-fold in *ΔS345* cells compared to WT (Figure 5B), consistent with the increased amounts of RoxS in the *ΔS345* strain. To confirm that the effect on the half-life of the *ppnkB* mRNA in this strain was due to the increase in RoxS levels, we constructed a strain lacking both sRNAs (S345 and RoxS). As expected, the *ppnKB* mRNA became stable again in the *ΔroxS ΔS345* double mutant with a half-life similar to a strain lacking RoxS alone (Figure 5B). This result confirms that the destabilization of the *ppnKB* mRNA in the *ΔS345* strain is RoxS-dependent. We thus propose that S345 be renamed RosA, for regulator <u>o</u>f <u>s</u>RNA <u>A</u>.

Intriguingly, unlike *ppnKB*, the rate of degradation of the *sucCD* mRNA was not affected in *ΔrosA* cells, with its half-life remaining at around 19 minutes in both the WT and the *ΔrosA* strain in LB medium (Figure 5B). We previously showed that RoxS is processed by RNase Y to remove the first 20 nts of the transcript producing a shorter, functional version of the sRNA called RoxS (D) ^14^. RoxS (D) is far more efficient at competing with the ribosome for binding to the *sucCD* transcript than the full-length RoxS sRNA *in vitro* ^14^. Removal of the first 20 nts of RoxS removes a significant portion of a 5’ stem loop, freeing up nucleotides to base-pair with the *sucCD* Shine-Dalgarno (SD) region. We therefore asked whether RosA had an effect on RNase Y processing of RoxS. RoxS (D) can be readily detected in a strain lacking the 5’-3’ exoribonuclease RNase J1 (encoded by *rnjA*), since this RNase is involved in the rapid degradation of the processed species. We therefore performed Northern blots on cells treated with rifampicin to compare the relative amounts and half-lives of RoxS and RoxS (D) in *ΔrnjA* versus *ΔrnjA ΔrosA* cells. Figure 6 shows that RoxS is efficiently processed to produce RoxS (D) in the *ΔrnjA* strain and is the most dominant form of RoxS in this strain. In contrast, in the double *ΔrnjA ΔrosA* mutant, full length RoxS was the dominant version of RoxS. Thus, RosA increases the efficiency of processing of RoxS to its truncated form. This is likely because base pairing with RosA is predicted to free up the RNase Y cleavage site in RoxS that is normally hidden within the duplex structure of the 5’ stem-loop (Figure 2A).

**Figure 6.**
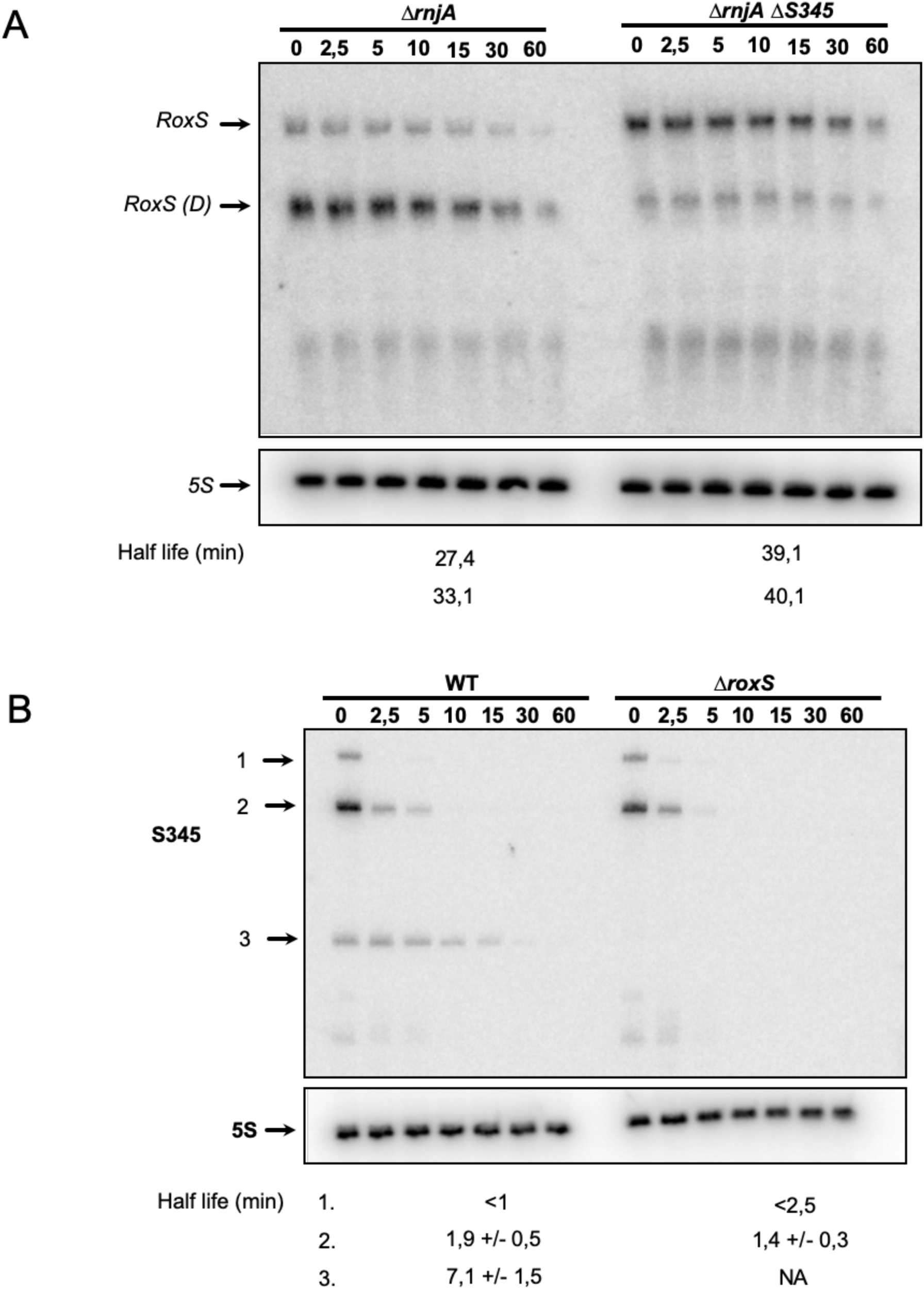
RNA processing events of S345 and RoxS are interdependent on the presence of the other. **A.** A representative Northern blot of total RNA isolated from *ΔrnjA* and *ΔrnjA ΔS345* strains probed for the RoxS sRNA at times after addition of rifampicin (Rif) at 150 μg/mL. The blot was re-probed for 5S rRNA as loading control. RoxS: Full size transcript, RoxS(D): truncated form of RoxS. Half-lives are given under each autoradiogram **B.** Northern blot of total RNA isolated from WT and *ΔroxS* strains probed for the S345 sRNA at times after addition of rifampicin (Rif) at 150 μg/mL. The blot was re-probed for 5S rRNA as loading control. The three forms of S345 are indicated by arrows. Calculated half-lives are shown beneath the autoradiographs and are the average of 2 experiments, with standard errors as shown.

To determine the global effect of RosA on RoxS and FsrA targets, and potentially identify other roles for this non-coding RNA, we performed a global proteomics analysis comparing the WT and *ΔrosA* deletion strains grown to mid-exponential phase in LB. The proteomes were analysed by label free quantitative proteomics. We detected 1463 proteins in the LC MS/MS analysis and identified 19 proteins that showed statistically significant (P value <0.05) reduced levels in the *ΔrosA* strain compared to WT (Table 1). Interestingly, seven of these proteins have already been assigned to the FsrA and RoxS regulons ^14, 15, 21^: CitB and SdhA have been assigned to the FsrA regulon, and PpnKB, CitZ, EtfA, SucC and SucD are members of the RoxS regulon. Most of the other proteins showing reduced levels in the *ΔrosA* mutant are predicted by CopraRNA or IntaRNA ^44, 45^ to be direct targets of FsrA and/or RoxS, and have been shown to bind similar metal ions and other cofactors to the proteins encoded by other RoxS/FsrA mRNA targets. This fits with the general agreement that members of the FsrA and RoxS regulons are involved in regulating genes involved in iron homeostasis and oxidoreduction ^14, 21^. The reduced levels of the FsrA and RoxS targets in the *ΔrosA* strain supports the idea that RosA counteracts regulation by both RoxS and FsrA and suggests that its primary role is as a sponge for these two sRNAs.

**Table 1.** Proteomics analysis for *ΔrosA*/WT shows reduced levels of RoxS and FsrA targets.

| Protein | BSU Number | log2(ΔS345/WT) | Copra/IntaRNA predicted target | Dysregulated in ΔFsrA or ΔRoxS | Protein description and Biological Process |
| --- | --- | --- | --- | --- | --- |
| AcsA | BSU29680 | -2.27 | RoxS |  | acetyl-CoA synthetase<br>utilization of acetate, fatty acids |
| SpoVS | BSU16980 | -1.57 | FsrA |  | Unknown<br>spore coat assembly, spore core dehydration |
| CitB | BSU18000 | -1.40 | FsrA | ΔFsrA | aconitase, trigger enzyme<br>TCA cycle |
| NadK2/<br>PpnKB<br>/YtdI | BSU29540 | -1.14 | RoxS | ΔRoxS | ATP-NAD kinase<br>NADP biosynthesis |
| YrhF | BSU27210 | -1.07 | RoxS |  | Unknown |
| EtfA | BSU28520 | -0.95 | RoxS and FsrA | ΔRoxS | electron transfer flavoprotein<br>(alpha subunit)<br>fatty acid degradation, |
| CitZ | BSU29140 | -0.95 | RoxS and FsrA | ΔRoxS | Citrate synthase 2<br>TCA Cycle |
| OdhB | BSU19360 | -0.87 |  |  | TCA Cycle<br>2-oxoglutarate dehydrogenase complex |
| OdhA | BSU19370 | -0.85 | RoxS |  | TCA Cycle<br>2-oxoglutarate dehydrogenase (E1 subunit) |
| SucC | BSU16090 | -0.84 | RoxS | ΔRoxS | succinyl-CoA synthetase (beta subunit)<br>TCA Cycle |
| YvyI/Pm<sub>i</sub> | BSU35790 | -0.78 | FsrA and RoxS |  | mannose-6-phosphate isomerase<br>mannose utilization |
| GudB | BSU22960 | -0.78 |  |  | glutamate dehydrogenase,<br>trigger enzyme<br>glutamate utilization, control of GluC activity |
| CitA | BSU09440 | -0.78 | FsrA and RoxS |  | minor citrate synthase<br>Unknown |
| SucD | BSU16100 | -0.76 | | ΔRoxS | succinyl-CoA synthetase (alpha subunit)<br>TCA Cycle |
| YpbR/<br>DynA | BSU22030 | -0.73 | RoxS |  | dynammin-like protein fusion of membranes |
| YkrA | BSU14550 | -0.62 | RoxS |  | Unknown |
| YcsA | BSU04000 | -0.62 | RoxS |  | putative tartrate dehydrogenase Unknown |
| SdhA | BSU28440 | -0.61 | FsrA | ΔFsrA | succinate dehydrogenase (flavoprotein subunit)<br>TCA cycle |
| YpiB | BSU22580 | -0.48 | FsrA |  | Unknown |

### Production of the short form of RosA requires RoxS

We showed above that RosA is important for the processing of RoxS to RoxS (D); we therefore wondered whether the converse was also true, *i.e.* whether RoxS had an effect on the processing of RosA. In Figure 6B we analysed the expression pattern and degradation rates of the three RosA species in the presence and absence of RoxS. In the presence of RoxS, all three forms of RosA were detected as seen above (Figure 4). However, in the absence of RoxS, the RosA species 3 was completely absent. We conclude that the interaction with RoxS plays a role in the processing of RosA to its smallest form. The size of species 3 (∼120 nts) is consistent with an RNA that extends from the mapped 5’ of all three RosA species to the end of the duplex with RoxS around nt 116 (Figure 2B). The duplex would protect the 3’ end of species 3 from 3’ exoribonucleases, consistent with its relatively long half-life compared to species 1 and 2. No effect on RosA processing could be seen upon deletion of FsrA under these growth conditions (Supplementary Figure 2).

### RosA provides a fitness benefit for *B. subtilis* under conditions of oxidative respiration

We asked whether RosA had an impact on cell doubling time by comparing the growth rate of the *ΔrosA* strain to that of the WT. No major difference in growth rate was seen in either LB or in M9. To ask whether there was a more subtle fitness cost to the cells lacking RosA, we performed competition assays between WT and *ΔrosA* cells in LB medium. We mixed the WT strain marked with a spectinomycin antibiotic resistance cassette and the phleomycin resistant *ΔrosA* mutant at a 1:1 ratio, which was confirmed by colony counts carried out on the starting culture. We then counted the number of *ΔrosA* and WT bacteria after 24 hours. The *ΔrosA* strain was recovered at significantly lower levels than the WT suggesting that it is at a competitive disadvantage (Figure 7). In a control experiment, we also competed a phleomycin resistant strain deleted for *yqbR*, a gene located on the Skin prophage region, that was shown to be transcriptionally inactive in LB by Nicolas et al ^12^. This strain retained a 1:1 ratio with the WT strain after 24 hours. We were also able to restore the fitness deficit of the *ΔrosA* strain with ectopic expression of RosA at the *amyE* locus. We propose that the reduction in the levels of enzymes of the TCA cycle, targeted by increased expression of FsrA and RoxS in the *ΔrosA* strain, gives these bacteria a fitness disadvantage as they are unable to generate ATP as quickly the WT strain.

**Figure 7.**
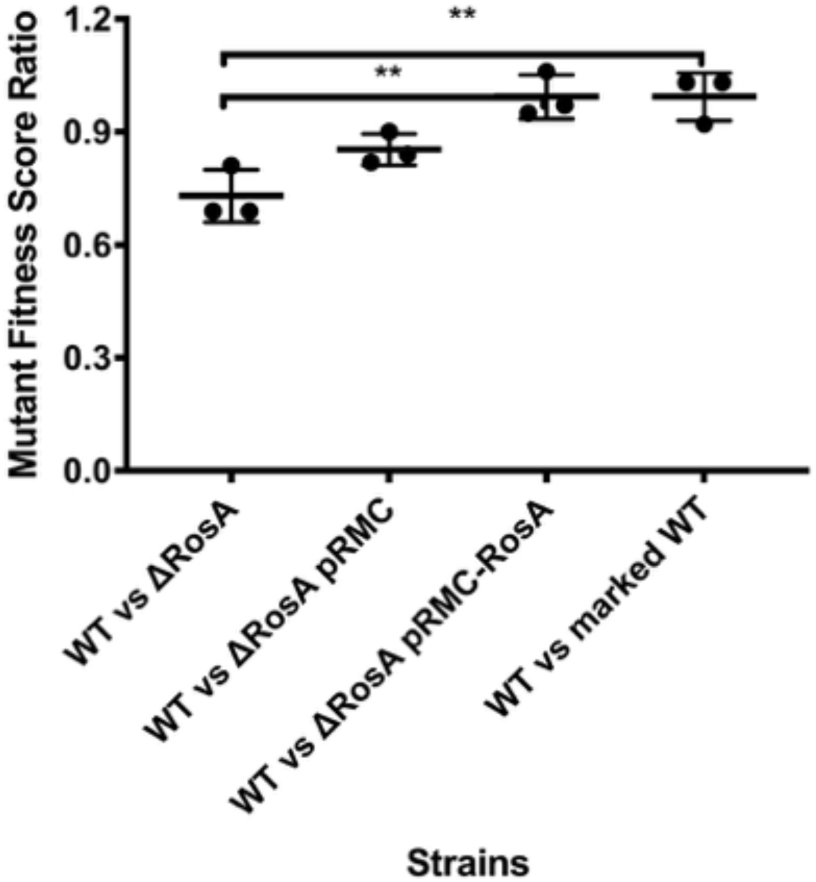
ΔS345 is reduced in fitness. The fitness deficit of the *ΔrosA* strain in LB was shown by co-culturing ΔS345 with the wild type strain mixed in a 1:1 ratio. The fitness deficit of *ΔrosA* was complemented by cloning the RosA sRNA under the control of its native promoter into the pRMC plasmid. Strains were grown for 24 hours and plated on antibiotics to enable CFUs to be determined for each strain in the mixed culture. An antibiotic marked wild type strain was used as a control. Statistically significant differences in fitness between strains calculated using Welch’s T test are shown by **. The experiment was repeated three times.

### RosA is subject to carbon catabolite repression

To begin to understand under which physiological conditions RosA might act as a sponge of FsrA and RoxS, we investigated how transcription of RosA is controlled. We used the DBTBS server to determine which transcription factors are predicted to bind to the RosA promoter region and regulate its transcription ^46^. DBTBS predicted a binding site for the transcriptional regulator CcpA between −1 to + 12 relative to the mapped 5’ end of RosA (Figure 1). CcpA mediates carbon catabolite repression in *B. subtilis*, repressing catabolic genes and activating genes involved in excretion of excess carbon ^47^. The prediction of a CcpA binding site in the promoter region of RosA was corroborated by Marciniak *et al.* who identified the coordinates of the CcpA binding site in front of RosA ^48^. The expression profile of RosA in the 104-condition tiling array data for *B. subtilis* was very similar to known members of the CcpA regulon, such as MalA, AcoA and AbnA, consistent with the idea that RosA is a CcpA regulated sRNA ^12^.

To confirm the regulation of *rosA* by CcpA we fused the promoter of *rosA* to GFP using the BaSysBioII vector ^32^. We monitored expression of this fusion in WT *B. subtilis* and in an isogenic mutant lacking the *ccpA* gene. No difference in P*rosA*-GFP expression could be seen between the WT and the Δ*ccpA* strain in LB medium (Figure 8A). Addition of 0.3 % (w/v) glucose to the medium resulted in repression of *rosA* promoter activity in the WT strain (Figure 8B), whereas in the absence of *ccpA* the *rosA* promoter remained active, as predicted (Figure 8B).

**Figure 8.**
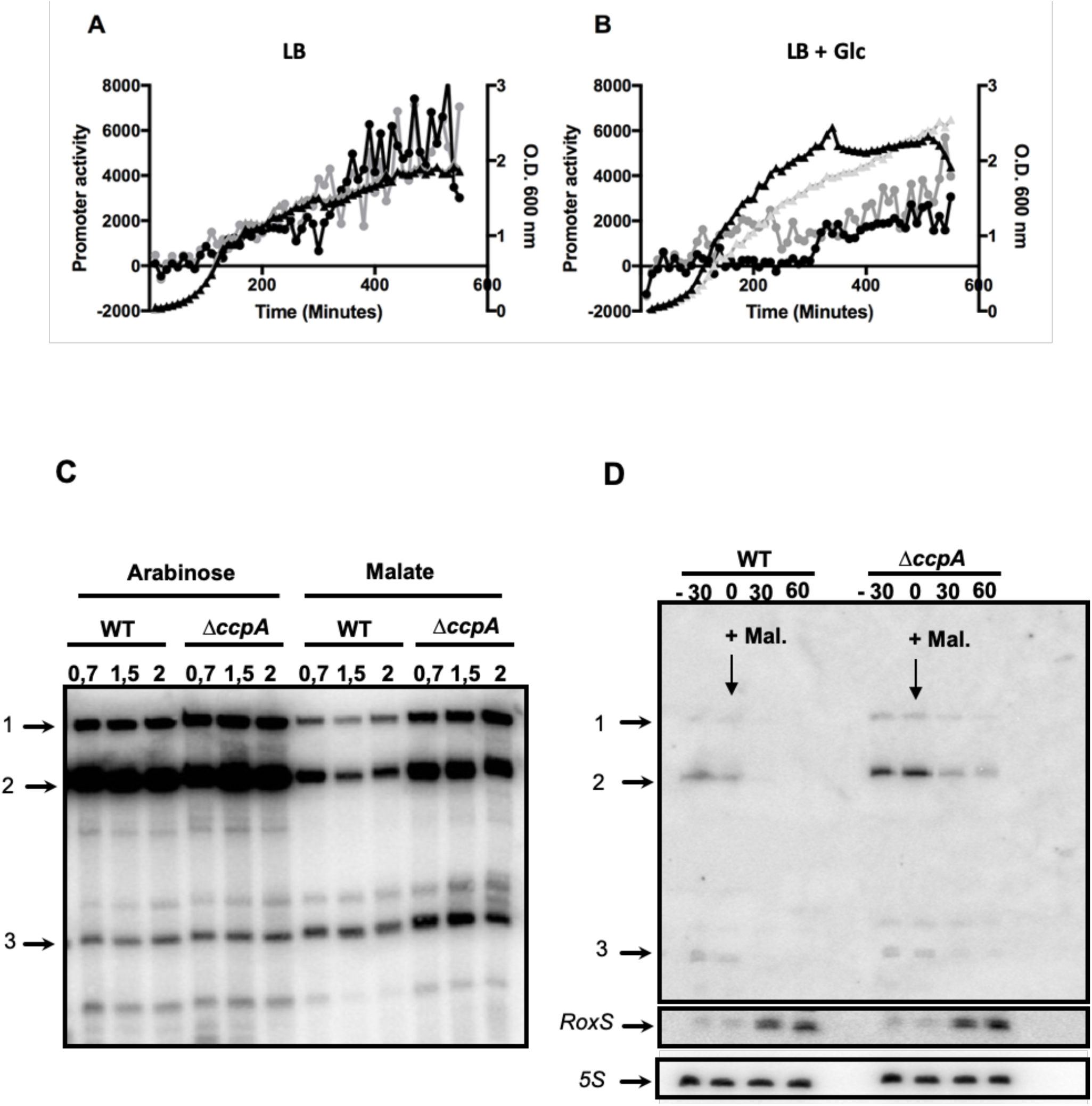
RosA is a CcpA regulated sRNA. **A.** Assay of the *RosA* promoter fused to GFP in wild type *B. subtilis* and Δ*ccpA* in LB with no glucose and **B.** with the addition of 0.3% glucose. Black line with triangles WT growth curve, grey line with triangles Δ*ccpA* growth curve. Black line with circles promoter activity of RosA in WT, grey line with circles promoter activity in Δ*ccpA*. Experiments were done in triplicate.**C.** Northern blot of RosA in WT and *ΔccpA* mutant strain grown in MD medium with arabinose (1%) or malate (1%) as carbon sources. RNA was extracted at several O.D._600_ during growth as indicated. **D.** Northern blot of RosA and RoxS in WT and *ΔccpA* mutant strain grown in MD medium with 1% arabinose. At mid exponential phase (O.D._600=0,6_), malate has been added to the culture and samples taken before (−30 min) and after (0, 30 and 60 min) the addition of malate. The blot was re-probed for 5S rRNA as loading control.

We performed a similar experiment where we measured the levels of RosA RNA in a defined medium with 1% malate or arabinose by Northern blot (Figure 8C). RosA levels were similar in WT and *ΔccpA* mutant strains grown in arabinose where CcpA is inactive on its targets. In contrast, as observed with the promoter fusion, RosA expression was repressed in the WT strain and this repression was alleviated in the *ΔccpA* mutant strain grown in a medium supplemented with malate.

We also measured RosA expression during a switch in carbon source. *B. subtilis* WT and *ΔccpA* strains were first grown to late exponential phase in a defined medium with arabinose as the sole carbon source, before adding 1% malate to promote carbon catabolite repression. Cells were harvested during exponential phase and 30 and 60 min after addition of malate and RosA RNA levels were measured by Northern blot (Figure 8D). Expression of RosA decreased after addition of malate in both the WT and *ΔccpA* strain. However, the level of RosA was higher in *ΔccpA* strain than in the WT demonstrating that RosA is subject to the catabolite repression, and that this regulation is partially CcpA dependent. In contrast, when the same membrane was re-probed for RoxS, we observed expression of RoxS was induced upon addition of malate as previously observed ^17^. These experiments confirm that RosA is a carbon catabolite responsive sRNA controlled by CcpA and that RoxS and RosA are important to manage the reprogramming of gene expression during a switch in carbon sources.

## DISCUSSION

In this study we report the use of *in vivo* RNA cross-linking using the psoralen AMT to globally identify RNA-RNA interactions occurring in the Gram-positive model organism *B. subtilis*. Our results identified hundreds of potential interactions, including previously well described sRNA-mRNA interactions. Two of three known sRNAs containing C-rich regions in *B. subtilis*, FsrA and RoxS, have been shown to target transcripts encoding essential components of central metabolism using their C-rich regions ^14, 15, 20^. In addition to the identification of known and new mRNA targets for RoxS and FsrA, we also showed that these two sRNAs interact with a new sRNA, S345, that we renamed RosA in this study.

Deletion of RosA from the genome of *B. subtilis* leads to a 3.5-fold increase of the half-life of RoxS showing that RosA controls RoxS turnover. In parallel, a proteomic analyses in the *ΔrosA* strain show a reduced levels of the known RoxS and FsrA targets like the TCA cycle enzymes, SucCD, OdhAB, CitZ, SdhA and CitB ^14, 15, 21^. Many of the other proteins with reduced levels, identified by the proteomic experiment, were also predicted to be targets of either RoxS or FsrA using CopraRNA ^44, 45^. A predicted target of RoxS is *acsA*, which encodes a key enzyme in central metabolism since it catalyses the conversion of ATP, acetate and CoA to AMP, diphosphate and acetyl-CoA, thus acting as a balancing point for the levels of CoA and acetyl-CoA in the cell ^49^. Furthermore, the SrtN protein is used by the cell to deacetylate AcsA and this reaction depends on NAD^+^ ^50^. The goal of RoxS-mediated reduction in AcsA levels may be to reduce non-essential NAD^+^ consumption. Moreover, we showed that the level of RosA and RoxS is comparable in LB and that one-to-one mixtures of RosA and RoxS *in vitro* result in full-duplex formation. Thus, these results show that RosA has the potential to be a highly efficient sponge of RoxS and FsrA activity in *B. subtilis* cells. Our data show that RosA acts differently on the two target RNAs. FsrA is sequestered in classic sponge RNA activity as the levels of the FrsA RNA remain unchanged, but its proteins targets are reduced in the absence of RosA. Whereas for RoxS we have shown that the levels, processing and its target efficacy are being affected.

In the field of eukaryotic RNA regulation, sponge RNAs are well-accepted as part of the regulatory landscape ^51^ and this idea has recently been getting increased traction in bacteria. Indeed, several sponge RNAs have been described in Gram-negative organisms and, intriguingly, many are derived from other transcripts (reviewed by Figueroa-Bossi and Bossi ^42^ and Azam and Vanderpool ^41^). In contrast, RosA is a stand-alone sRNA. Interestingly, another stand-alone sRNA in *S. aureus*, namely RsaI (RsaOG), was also shown to be CcpA-regulated and to interact with the sRNAs RsaG, RsaD and RsaE, the RoxS homologue in *S. aureus*, ^52^. RsaI, like RosA, contains two G-rich regions to bind to RsaG, RsaD and RsaE. These results suggest that RsaI and RosA could fulfil the same functions in *S. aureus* and *B. subtilis* and that similar sponge RNA-mediated regulatory pathways exist in Firmicutes to balance the metabolic requirements of the cell. Indeed, RsaI is conserved in the genus *Staphylococcus* but not in Bacilli, while RosA is conserved in some Bacilli but not in the Staphylococci. The role of RsaI as a sponge RNA remains to be definitively proven since the impact of RsaI on RsaE, RsaD and RsaG mRNA targets has not yet been investigated. In contrast to RosA, RsaI has also been shown to additionally have a C-rich region used to bind mRNA targets. It could thus act as both a direct regulator and as an sRNA sponge. The absence of equivalent C-rich regions in RosA may limit its function to that of a sponge RNA. However, we plan to study whether RosA can directly regulate its own mRNA targets.

In this study, we identified three forms of RosA, with different half-lives. Full length RosA (229 nts) is very short-lived and is quickly processed to the 185 nts form that appears to be the main functional form of RosA. The shortest form of RosA (species 3) has a half-life of 7.1 minutes and its generation is RoxS dependent. We believe that this form of RosA corresponds to a stable degradation product protected from 3’ degradation by duplex formation with RoxS. None of the three most commonly used RNases in *B. subtilis*, RNase J1, RNase III or RNase Y could account for the processing of RosA to its different forms (data not shown). The role of RosA in facilitating the processing of RoxS and the possible persistence of a RoxS-RosA duplex in cells (RosA species 3) raises the interesting question of whether RoxS can be recycled from RosA to regulate mRNAs such as *sucCD* that prefer the shorter form of RoxS? One could imagine that this duplex might be a reservoir of mostly processed RoxS, that could switch to new partners for which it had a greater affinity. Further experiments are required to explore this possibility.

Expression of RoxS is tightly controlled by two transcription factors, ResDE and Rex ^14, 17^. Why then is this additional level of post-transcriptional regulation of RoxS by RosA required? Our previous data suggests that RoxS is involved in readjusting the transitory imbalances in NAD/NADH ratio that occur upon encountering carbon sources such as malate. Through its role in reducing NADH levels, RoxS eventually increases the DNA binding capacity of the transcriptional activator Rex, turning down its own expression. However, RoxS is a relatively stable sRNA, with a half-life of 13 min in a WT strain that increases to >45 min in the absence of RosA. The use of this non-coding sponge RNA is thus likely be a way to dial down RoxS activity more efficiently than by simply turning off transcription, first by neutralizing the C-rich regions involved in the regulation of all known targets so far and then by stimulating its degradation. We propose that RosA accelerates the degradation of RoxS by stimulating the opening of the 5’ stem loop of RoxS, where RNase Y is known to cleave to produce the truncated form of RoxS, named RoxS (D), *i.e.* the processing pathway that leads to the functional form of RoxS required for *sucCD* regulation is also the first step in RoxS turn-over. In agreement with this hypothesis, the 45 min half-life of RoxS observed in the *ΔrosA* strain is similar to that measured previously in a strain deleted for RNase Y ^14^.

We determined that RosA is transcriptionally repressed by the main carbon catabolite repressor in *B. subtilis*, CcpA, and that the expression of RoxS and RosA is anticorrelated during a switch of carbon source. When *B. subtilis* is grown on one of its preferred carbon sources such as malate ^53^, a large proportion of the carbon is metabolized only as far as pyruvate and acetyl CoA by malate dehydrogenase (Figure 9). These enzymes use NAD as a co-factor, leading to an increase of NADH concentration in the cell known to inhibit the DNA binding abilities of the transcriptional regulator Rex. This inhibition allows the transcriptional derepression of RoxS and, instead of directing malate into the TCA cycle, malate is converted to lactate and acetate *via* fermentation pathways normally repressed by Rex. Fermentation allows the regeneration of NAD+ from NADH. Concomitantly, RosA is repressed by CcpA allowing RoxS to bind its targets including mRNAs encoding enzymes of the TCA cycle which use NAD as co-factor. CcpA enables *B. subtilis* to quickly adapt to the presence of these preferred carbon sources. Indeed, CcpA represses genes involved in the metabolism of secondary carbon sources and turns down expression many of the enzymes of the TCA cycle and transporters of TCA cycle-intermediates, to ensure resources are not wasted ^47, 54^ (Figure 9). CcpA also activates the transcription of genes whose products are responsible for overflow metabolism when the bacteria are grown on a preferred carbon source. The targeting of these metabolic pathways is strikingly similar to what was observed previously by Durand *et al.* for RoxS, *i.e.* CcpA and RoxS have many overlapping targets ^17^ (Figure 10). In contrast, when *B. subtilis* is grown on a non-preferred carbon sources like arabinose, the inactivation of the carbon catabolite protein CcpA will allow the transcriptional derepression of the RosA sRNA and other CcpA regulated genes, including those encoding enzymes of the TCA cycle. RosA in turn sponges RoxS and impairs the post-transcriptional repression of RoxS targets, also including mRNAs implicated in the TCA cycle. Rex, for its part, represses the fermentation pathways (Figure 10).

**Figure 9.**
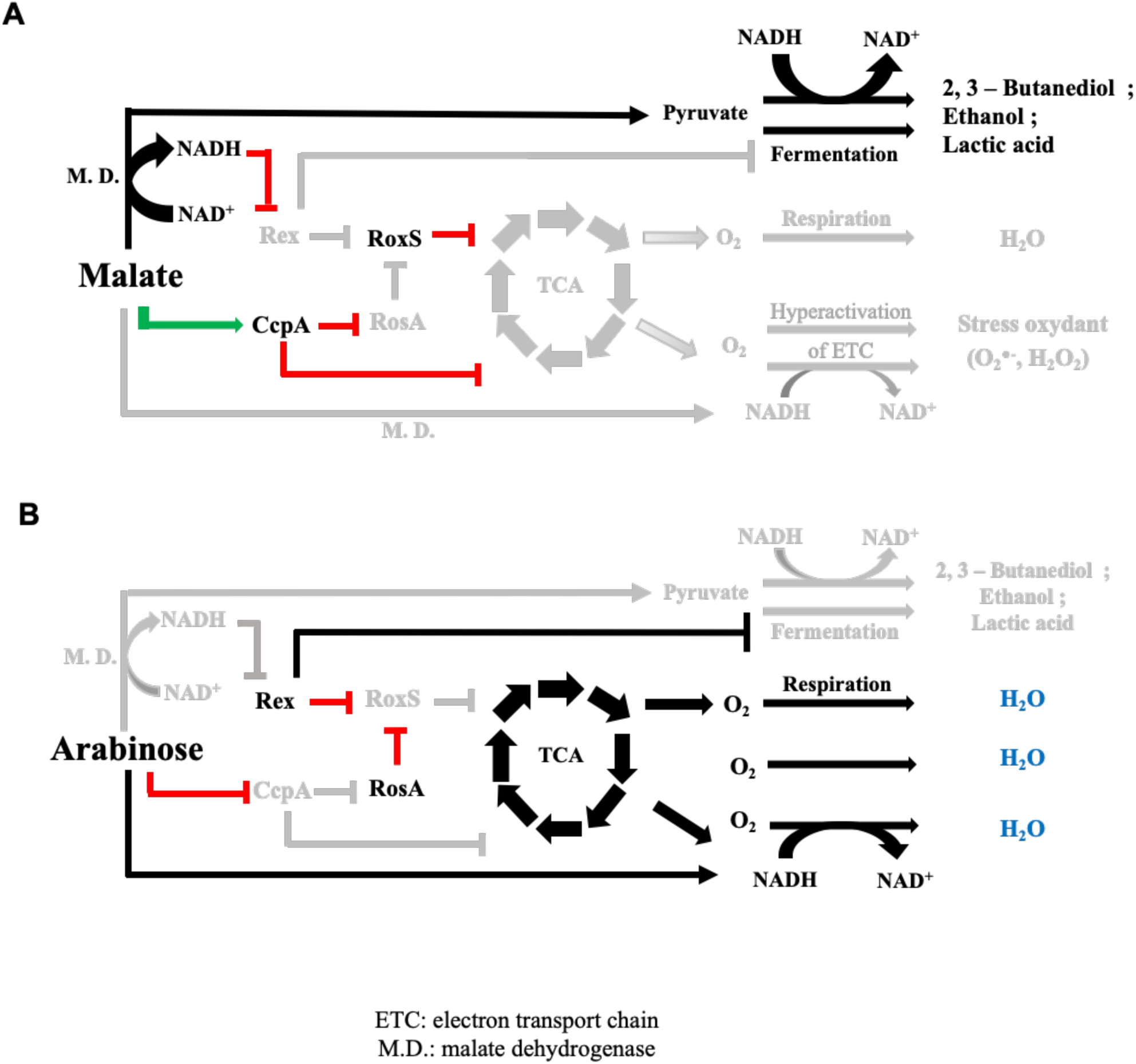
Model for the regulation of the fermentation and respiration pathways by the two sRNAs RoxS and RosA. **A.** In the presence of the preferred carbon source malate, malate dehydrogenases (M.D.) convert malate into pyruvate using NAD as co-factor. The increase of the NADH pool in the cell leads to the inhibition of Rex activity. RoxS is derepressed and regulates its targets (including TCA cycle and respiration enzymes). CcpA is activated and represses numerous genes including those encoding enzymes of the TCA cycle and RosA sRNA avoiding its sponging effect on RoxS. The goal of this regulation is dual: 1) To avoid the hyperactivation of electron transport chain (ETC) due to the increase of the NADH pool to limit oxidative stress 2) To activate fermentation pathways in response to the Rex inhibition to regenerate NAD. **B.** In the presence of a non-preferred carbon sources like arabinose, the high NAD/NADH ratio activates Rex which in turn represses fermentation pathways and RoxS sRNA. The carbon catabolite control by CcpA is also inhibited, allowing RosA expression. RosA sponges RoxS sRNA present in the cell and blocks its activity. This cascade of regulation allows the full activation of the TCA cycle and the aerobic respiration of the cell.

The discovery here of the RosA RNA sponge under the control of the transcription factor CcpA, provides the missing link between RoxS and CcpA. In other words, RoxS is connected to the CcpA regulon *via* the RosA non-coding RNA, and RoxS ensures an additional, potentially more rapid control at the post-transcriptional level for more than 30 % of genes that are regulated by CcpA. The effect of RosA on RoxS also significantly expands CcpA regulon.

## CONCLUSIONS

We have shown that *in vivo* AMT crosslinking of RNA is a suitable method to identify novel RNA-RNA interactions including sRNA interactions. We have focused here on a novel interaction between the two sRNAs FsrA and RoxS with the RNA sponge S345 that we have renamed RosA (Regulator of sRNA A) and have highlighted its role in balancing the metabolic state of the cell. However, there remains many newly identified interactions in the interaction data set to be investigated that likely represent many novel regulatory mechanisms.

## AUTHOR CONTRIBUTIONS

Conceptualization, ELD; Methodology, ELD, ACS, JRHF, CC^1^, SD, AM and TA; Investigation, ELD, SD JM, ACS, SL, GK and JRHF Writing – Original Draft, ELD, SD and CC^2^; Funding Acquisition, ELD, CC^1^ and SD; Supervision, ELD, CC^1^, TA, SD and AM.

## SUPPLEMENTARY DATA

Supplementary Data are available.

## ACKNOWLEDGEMENTS

The authors would like to thank Jan Maarten van Dijl and Ruben Mars for providing strains. Thomas Guest for constructing the RosA promoter fusion, and Justin Merritt and Holly Hall for useful discussion.

## Supplemental Data

**Supplementary Figure 1.**
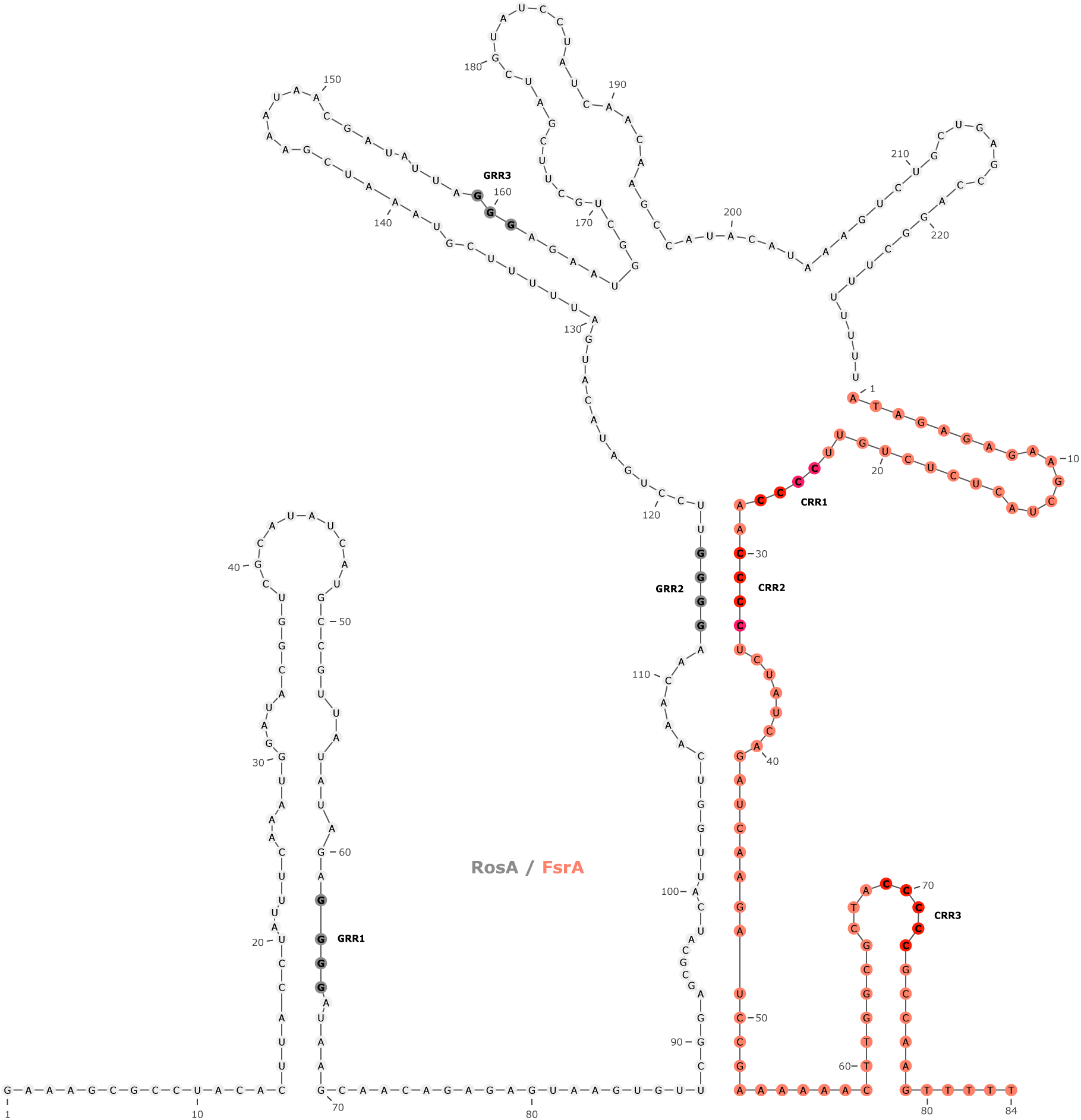
Prediction of the interaction between S345/RosA and FsrA. The interaction between FsrA and S345/RosA sRNAs was predicted with IntaRNA web server (http://rna.informatik.uni-freiburg.de/IntaRNA/Input.jsp). FsrA sRNA is coloured in orange. The C-rich region of FsrA and the G-rich region of S345 are coloured in red and grey respectively.

**Supplementary Figure 2.**
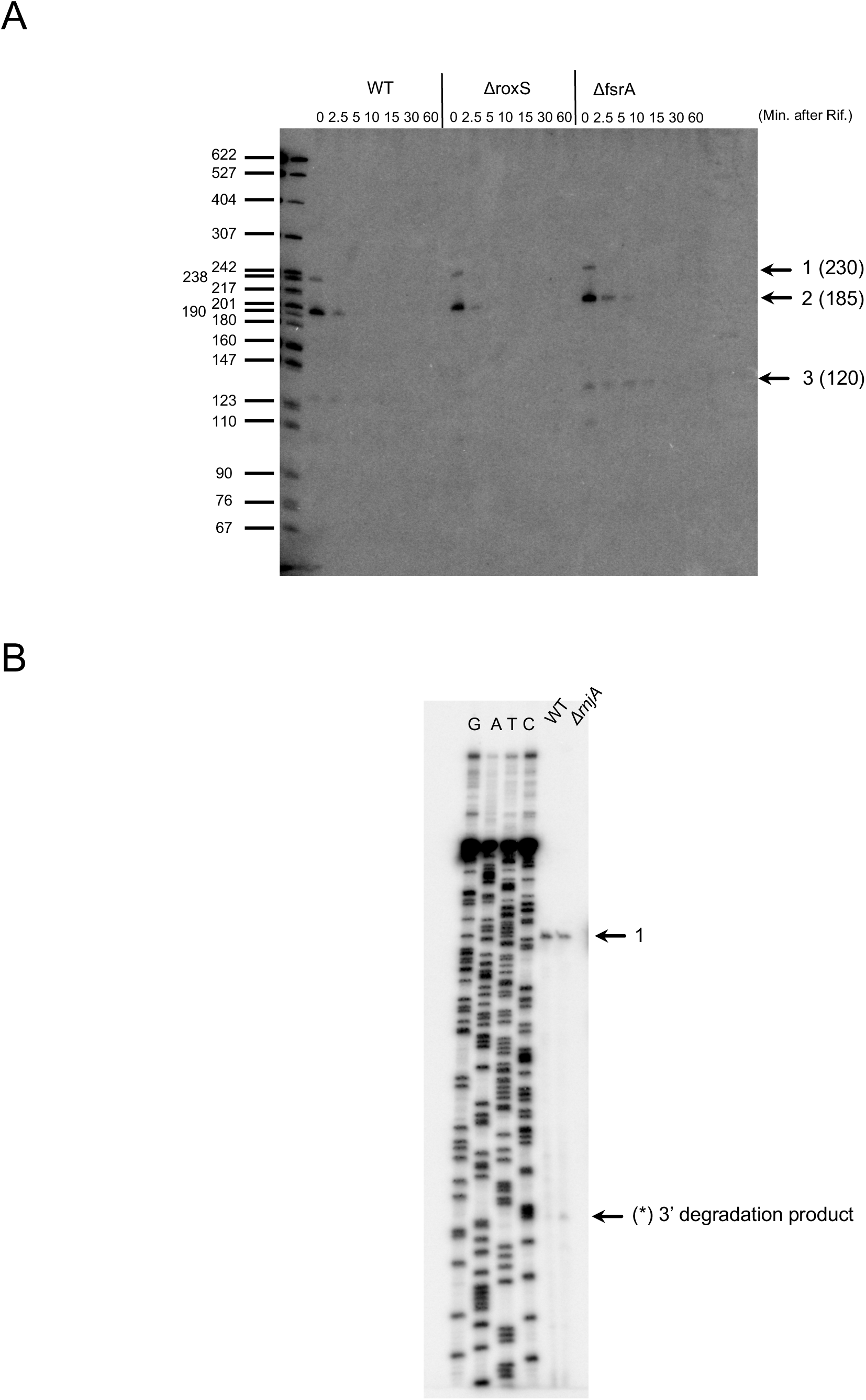
Sizes of S345. **A.** Northern blot analysis of S345/RosA in the WT, *ΔroxS* and *ΔfsrA* strains before and after the addition of rifampicin to inhibit transcription. The three forms of S345/RosA are indicated by arrows. A radiolabelled DNA marker was loaded on the left side. **B.** Primer extension to determine the 5’-end of S345. RNA was extracted from WT and *ΔrnjA* strains grown in LB. The sequencing reaction and the primer extension was done with an oligonucleotide closed to the 3’ end of S345/RosA. (1) indicates the 5’ end of S345. (*) The 5’ end of a 3’ degradation product.

**Supplementary Figure 3.**
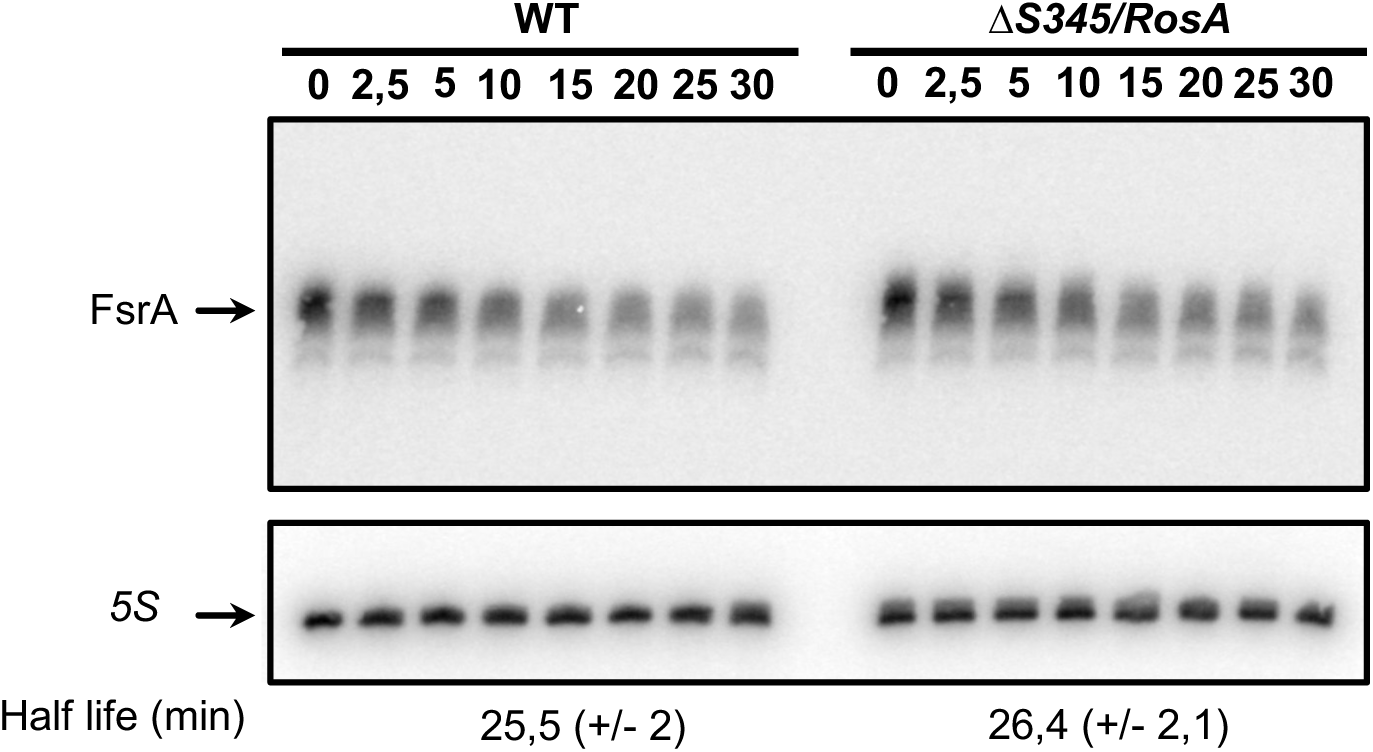
S345/RosA has no reciprocal impact on FsrA degradation or processing. Northern blot analysis of the FsrA sRNA in WT and *ΔS345/RosA* strains before and after the addition of rifampicin to inhibit transcription. Calculated half-lives are shown beneath the autoradiographs and are the average of 2 experiments, with standard errors as shown.

**Supplementary Figure 4.**
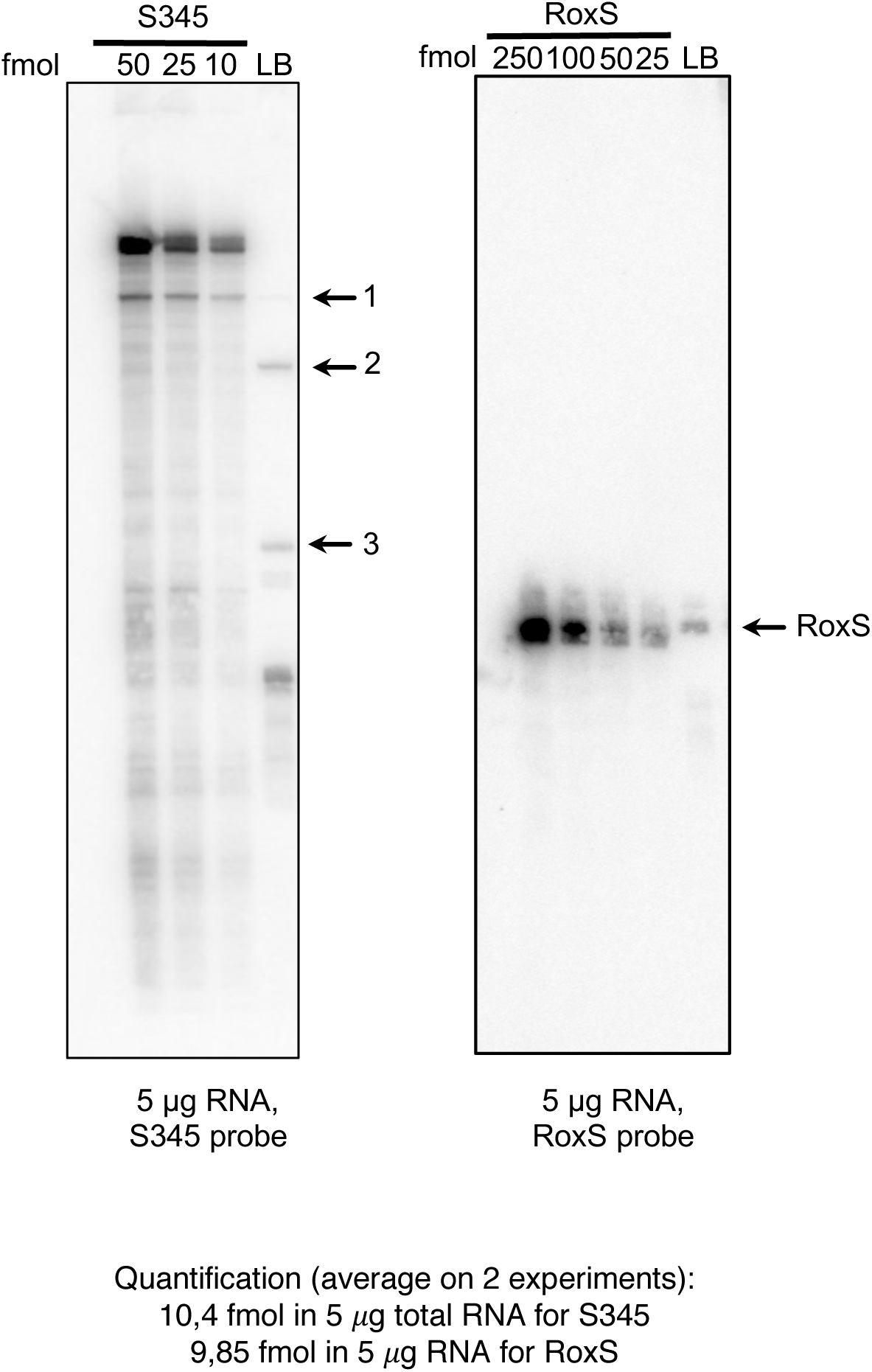
Quantification of S345/RosA and RoxS. Northern blot with a defined amount of *in vitro* transcript of S345/RosA (left side) and RoxS (right side) and 5 *μ*g of total RNA extracted from a WT strain grown in LB. The experiment was repeated 2 times.

**Supplementary Tables 1 and 2**

Strains and Oligos used in this study

**Supplementary Table 3**

All statistically significant interactions between FsrA (A) and RoxS (B). Columns - Sample (condition and strain), Target (interacting feature name), Target id (BSU number, Nicolas et al locus id, UA id), P value, sRNA_target_interaction (count of sRNA interactions with target), other_sRNA_interaction (count of other sRNA interactions in individual sample), other_target_interaction (count of other target interactions in individual sample), total_sRNA_reads (total read count for sRNA in sample, interacting and non-interacting), total_target_read (total read count for target in sample, interacting and non-interacting).

**Supplementary Table 4**

Most statistically significant interactions between FsrA (A) and RoxS (B) across the pairs of sequenced samples.

